## Supplementary figures and legends for version2 for "14-3-3 protein augments the protein stability of phosphorylated spastin and promotes the recovery of spinal cord injury through its agonist intervention": Supplementary figure legends.docx

**Figure S1. The peptides of 14-3-3 essential for the interaction with spastin are highly conserved in all 14-3-3 isoforms.** (A) The peak chart of binding peptides from LC-MS to spastin were shown. (B) Different 14-3-3 protein isoforms (rat species) were aligned and the LC-MS peptides were cropped to show the conservatism.

**Figure S2. The phosphorylation of Ser233 in spastin does not affect the microtubule severing activity of spastin.** GFP and GFP tagged spastin or its S233 phosphorylation mutants (S223A and S233D) were transfected into the COS7 cells. Cells were then fixed and stained with tubulin antibody to visualize the microtubule. Both GFP tagged spastin S233A and spastin S233D could sever the microtubules efficiently.

**Figure S3. 14-3-3 regulates spastin-dependent microtubule severing ability.** (A) GFP tagged spastin were transfected with flag tagged ubiquitin into COS7 cells, and cells were stained with β-tubulin antibody. (B) Under the same conditions, mCherry tagged spastin were transfected together with different Ser233 spastin mutations (spastin S233A or spastin S233D). Cells were also stained with β-tubulin to visualize the MT cytoskeleton. Scar bar, 50 μm and 10 μm.

**Figure S4. 14-3-3 agonist FC-A promoted neuritie outgrowth of hippocampal neurons in stage 2-3 via spastin.** (A) Hippocampal neurons at DIV2 were transfected with GFP plasmids together with or without interference to knockdown spastin expression, to visualize the morphology of neuron. Neurons were then incubated with FC-A for 24 hours, then cells were fixed and images were taken by confocal microscopy. Neurons were traced using Image J Pro Plus and quantitative analysis of the length of total neurites (B) and (C) the number of total neurites. **P* < 0.05, ***P* < 0.01, ****P* < 0.01. Scale bar: 100 μm.

**Figure S5. 14-3-3 inhibitor R18 inhibited neurite outgrowth of hippocampal neurons in stage 2-3 via spastin. (A)** Hippocampal neurons at DIV2 were transfected with GFP or GFP tagged spastin, together with R18 or not accordingly. Cells were fixed and images were taken by confocal microscopy. Neurons were traced using Image J Pro Plus and quantitative analysis of (B) the length of total neurites and (C) the number of total neurites. **P* < 0.05, ***P* < 0.01, ****P* < 0.01. Scale bar: 100 μm.

**Figure S6. 14-3-3 agonist FC-A promoted neuritie regeneration of cortical neurons in via spastin.** (A) Cortical neurons were cultured for 7 days and subjected to mechanically scratched with a pipet tip. Cells were incubated with FC-A or spastazoline for 24 hours. Cells were fixed and stained with βIII-Tubulin. Scare bar: 100 μm. (B) Quantification of the longest regenerative neurites. **P* < 0.05, ***P* < 0.01, ****P* < 0.01. Scale bar: 100 μm.

**Figure S7.** After spinal cord contusion, the 14-3-3 agonist FC-A and spastin inhibitor spastazoline were administered. Hematoxylin and eosin (H&E) and Luxol fast blue stain (LFB) were used for staining. Demyelination of the spinal cord can be observed within the region enclosed by the dotted line. This image is an enlargement of Figure 5J. Scale bar，2000 μm.

**Figure S8.** Following spinal cord contusion, 14-3-3 agonist FC-A and spastin inhibitor spastazoline were administered. Sections were stained with NF (neurofilament) and 5-HT antibodies. This image corresponds to Figure 5L and is an enlargement of it. The white box outlines the regions figure 5L1-L4 from the original image. Scale bar, 400 μm.

**Figure S9. Microtubule stability of in the lesion site of mice after spinal cord contusion and administrated with FC-A and spastazoline.** (A)The sagittal overview of the lesion site of spinal cord injury. Acetylated tubulin (green) and β-tubulin (red) were stained to label stable MTs and total MTs. Scale bar, 200 μm and 50 μm. (B) The intensity and proportion of acetylated tubulin and β-tubulin around the lession site in different groups (sham, injury, FC-A, spastazoline and FC-A+spastazoline). Scale bar, 50 μm. (C) Higher resolution image in panel B.

**Figure S10.** **Microtubule stability of in the lesion site of mice in a T10 lateral hemisection spinal cord injury model after relative drug administration.** (A) Sagittal view of acetylated tubulin (green) and β-tubulin (red) immunofluorescence in the white manner of the lesion site after SCI. Scale bar, 50 μm. (B) Quantification of the ratio of acetylated tubulin to β-tubulin immunoreactive fluorescence intensity (0.5 mm caudal to the lesion site).

**Figure S11. 14-3-3 agonist FC-A promotes the Nestin expression which was NeuN-positive. (A)** Sagittal view of Nestin (green) and NeuN (red) immunofluorescence in the white manner of the lesion site after SCI. Scale bar, 50 μm. (B) Quantification of the ratio of Nestin to NeuN immunoreactive fluorescence intensity (0.5 mm caudal to the lesion site).

**Figure S12. 14-3-3 agonist FC-A promotes the Nestin expression which mainly co-localized with Brdu.** Sagittal view of Brdu (green) and Nestin (red) immunofluorescence in the white manner of the lesion site after SCI. Scale bar, 500 μm.

**Figure S13. Footprint assay of mice after the intervention by FC-A and spastazoline following spinal cord injury.** Example images depicting footprint analysis on 16 DPI (days post injury). The measurement of the stride length and width were shown in red line. Scale bar: 2.5 cm.
