## Supplementary figures and images for "14-3-3 protein augments the protein stability of phosphorylated spastin and promotes the recovery of spinal cord injury through its agonist intervention"

### S1.tif

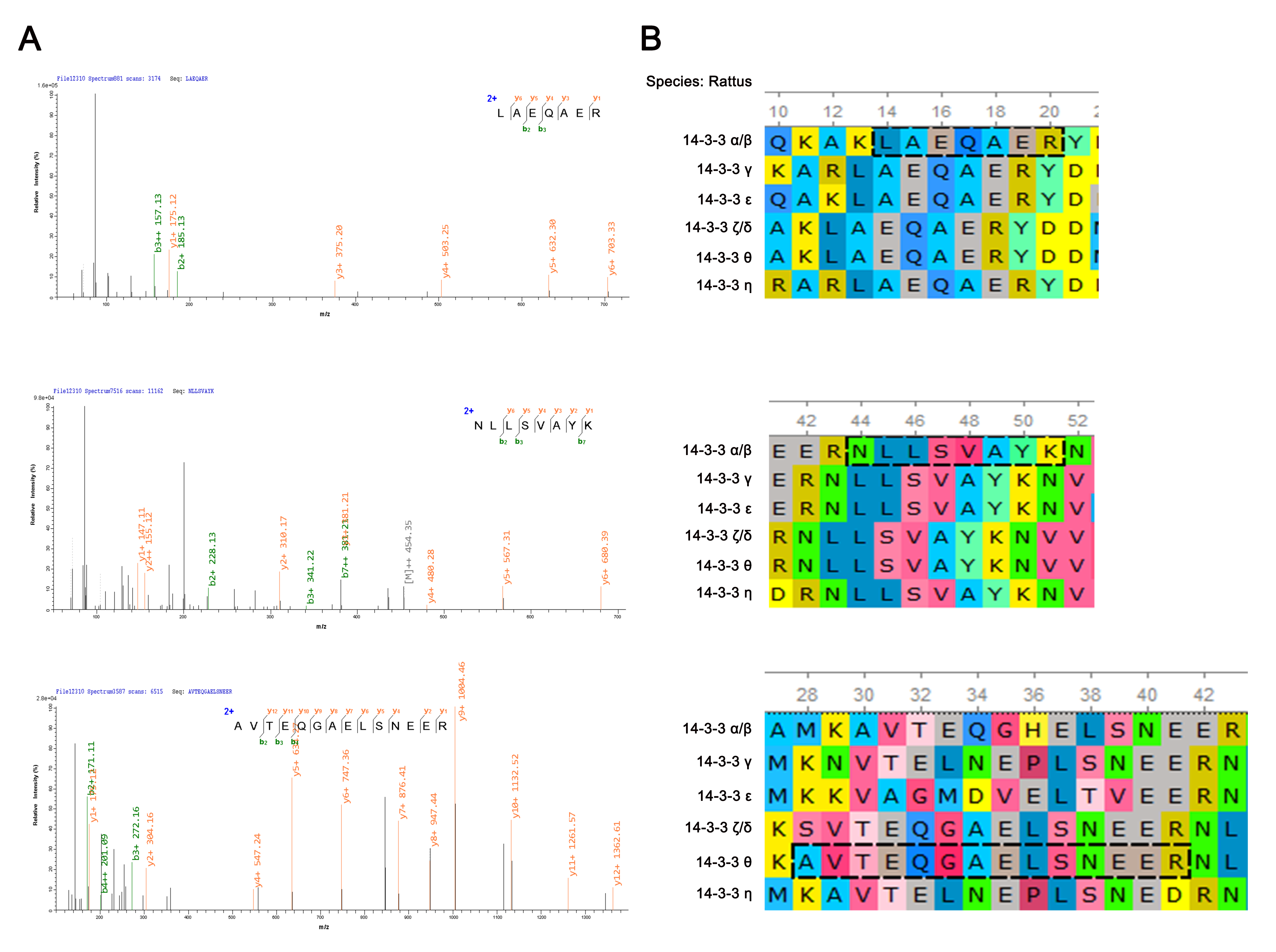

### S2.tif

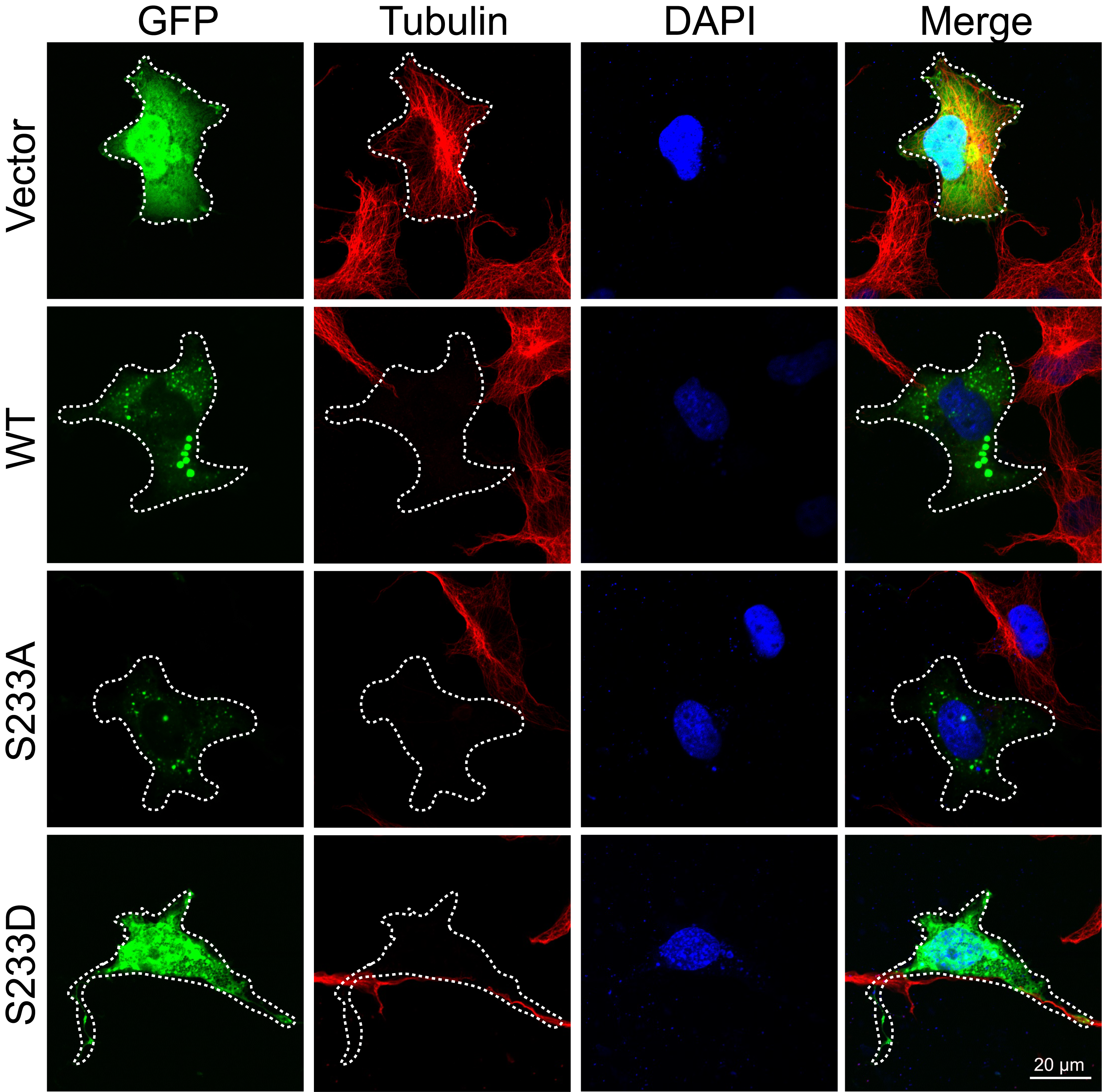

### S3.tif

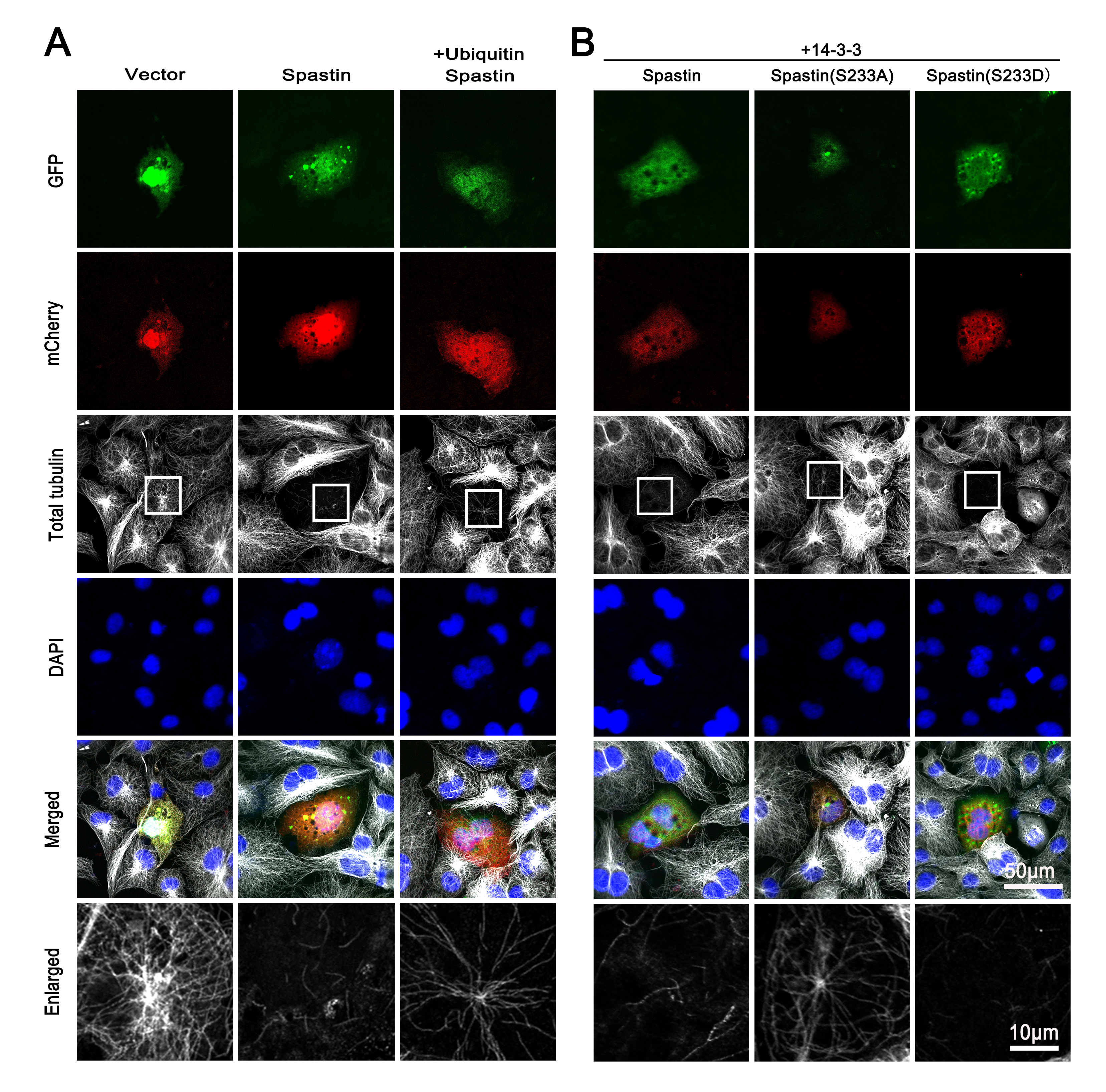

### S4.tif

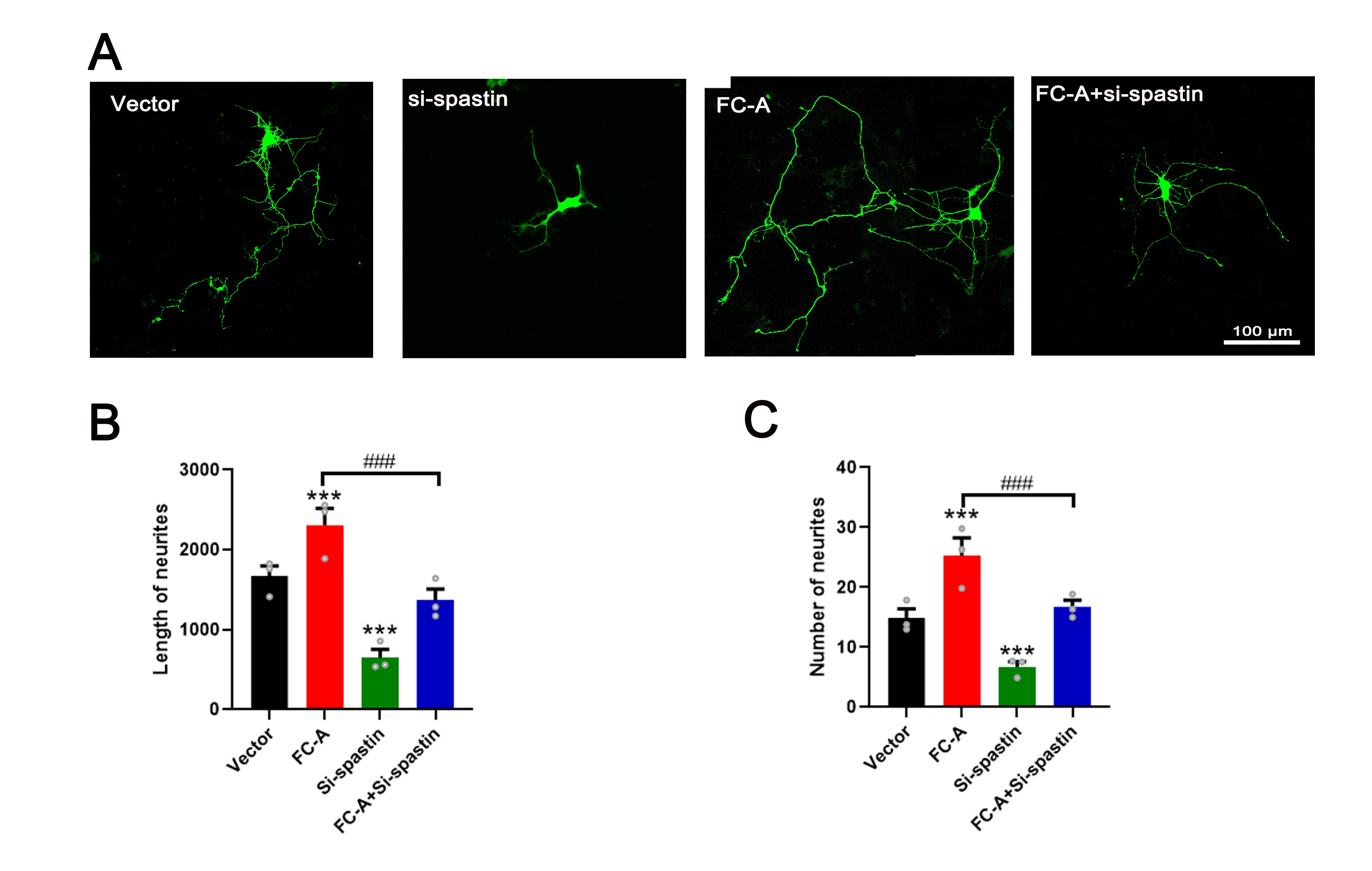

### S5.tif

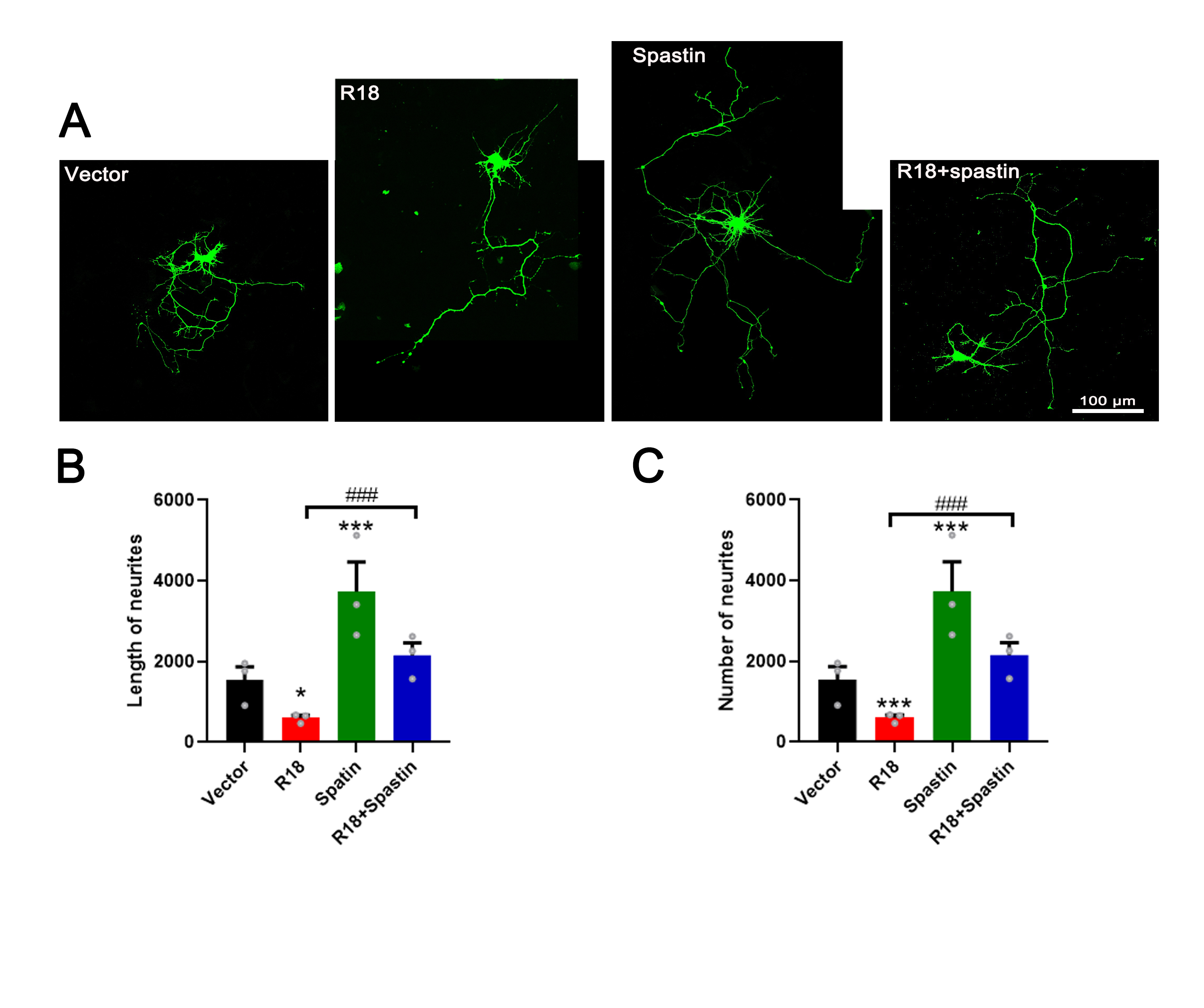

### S6.tif

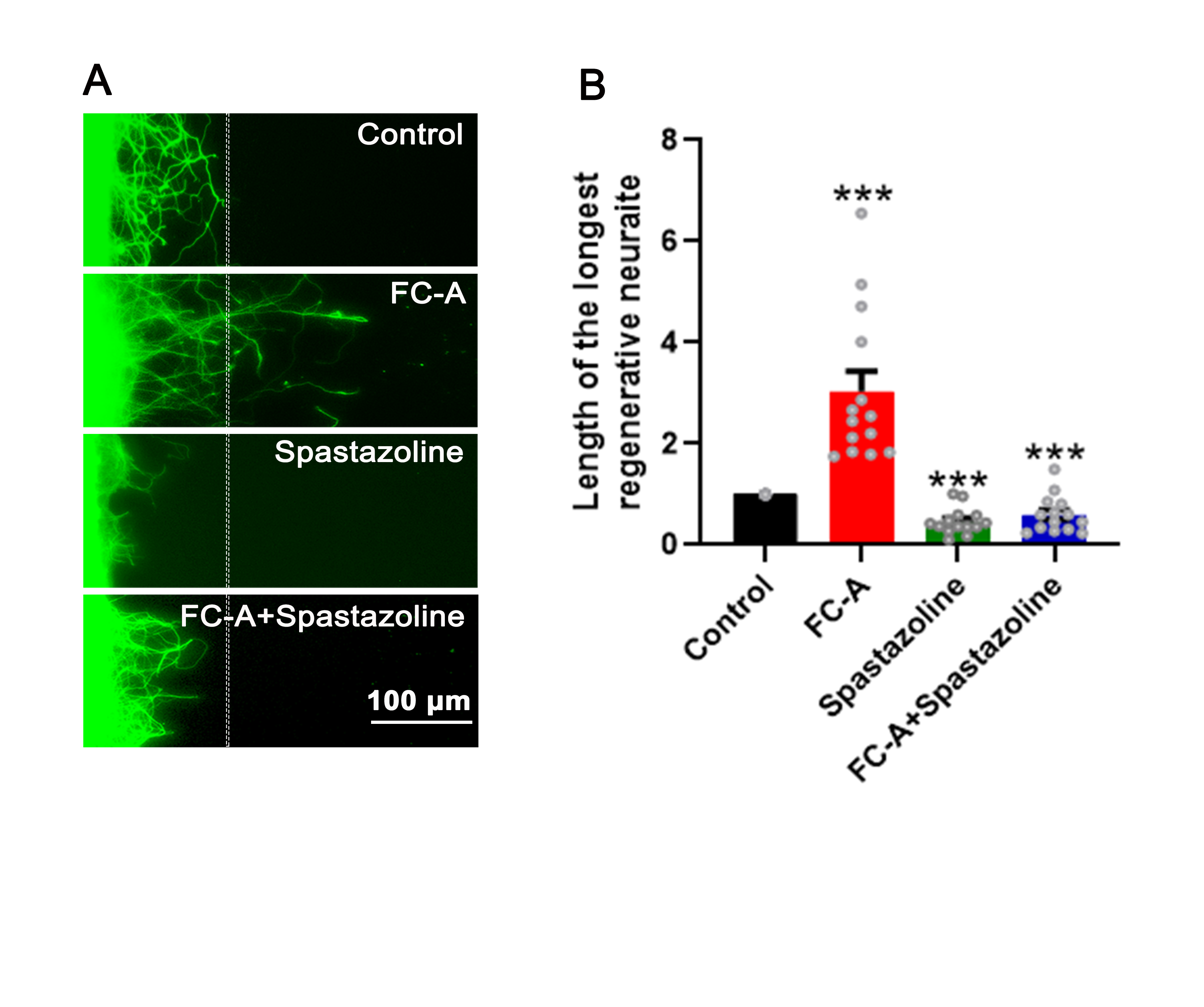

### S7.jpg

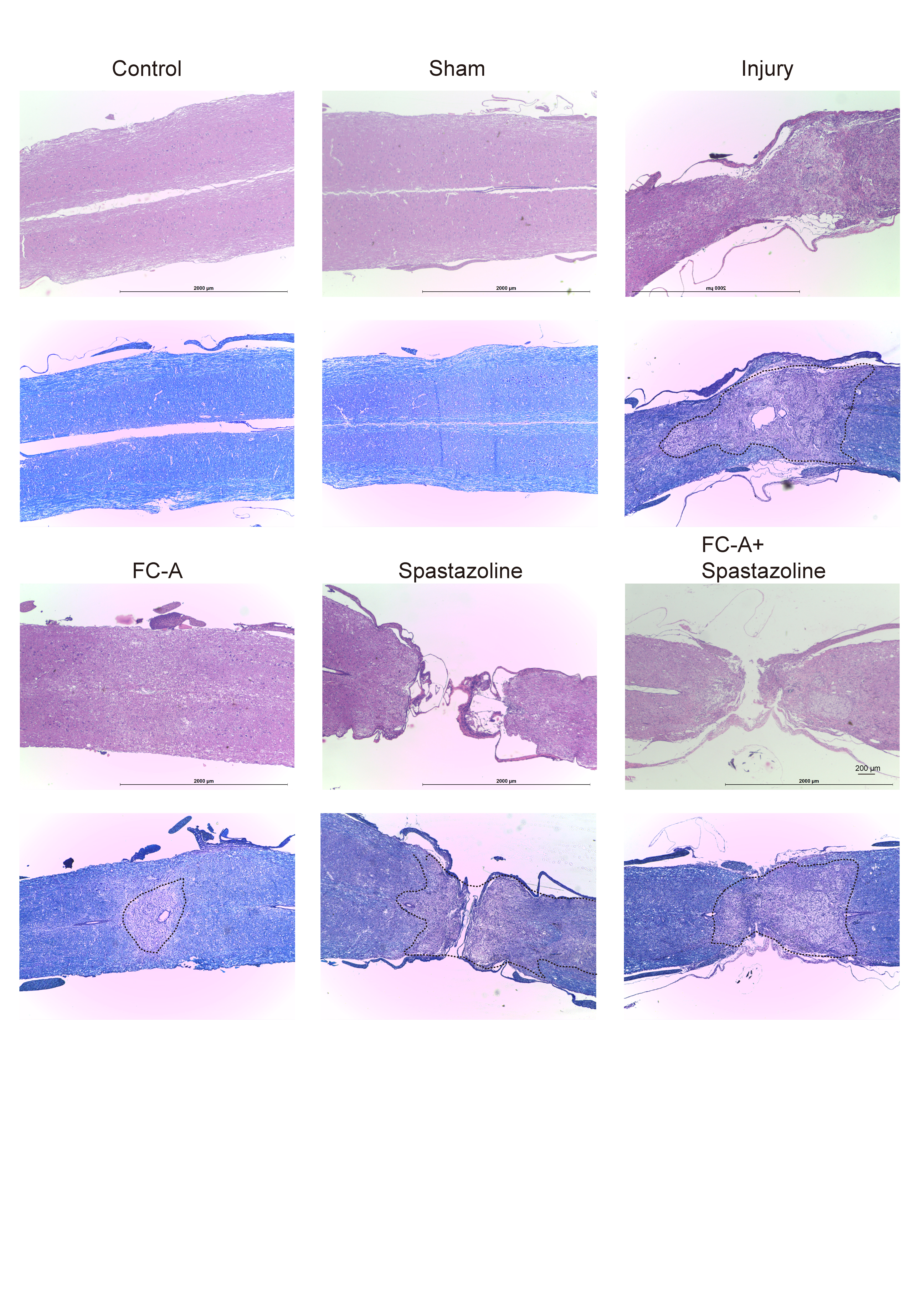

### S8.jpg

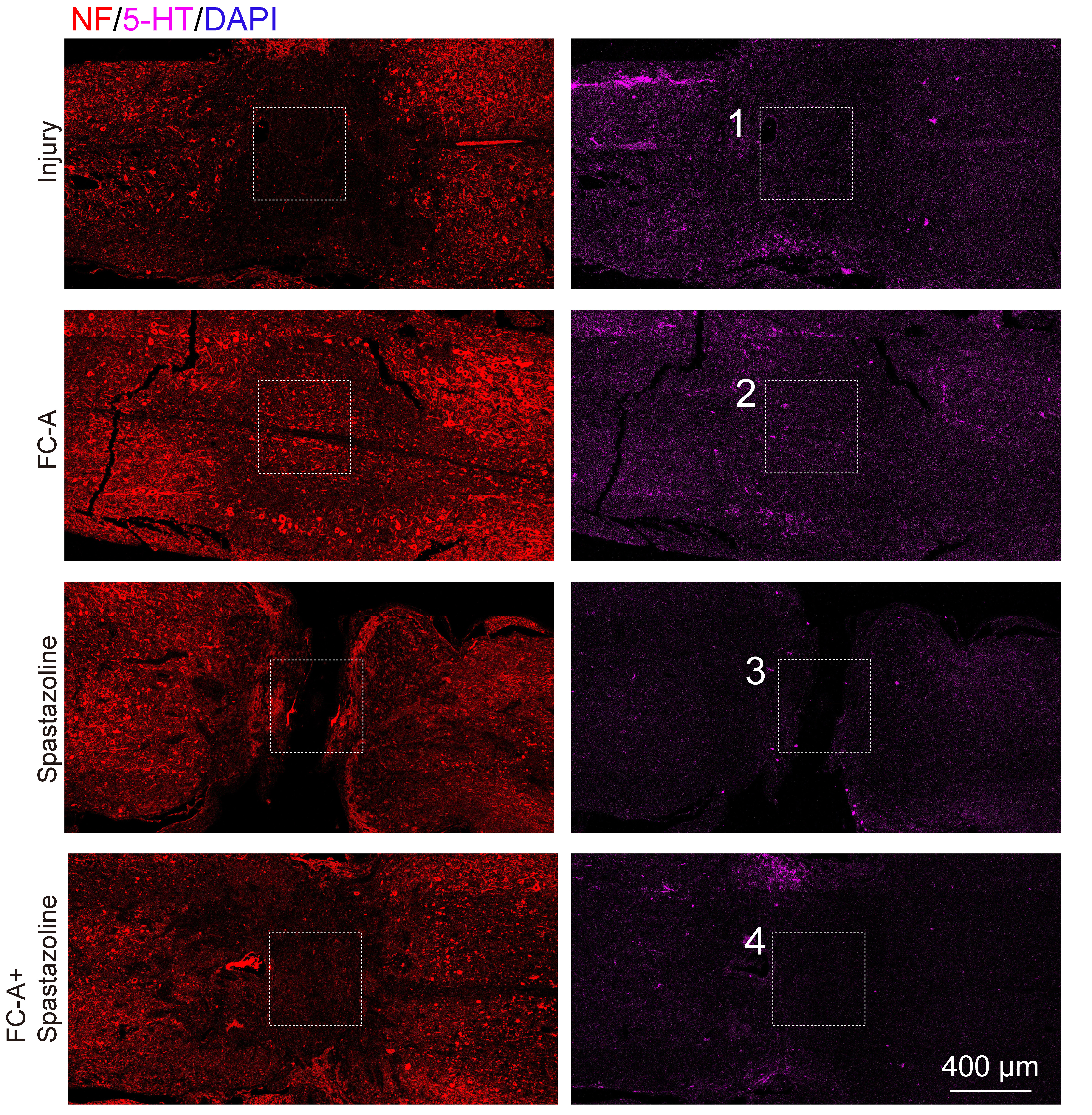

### S9A&B.tif

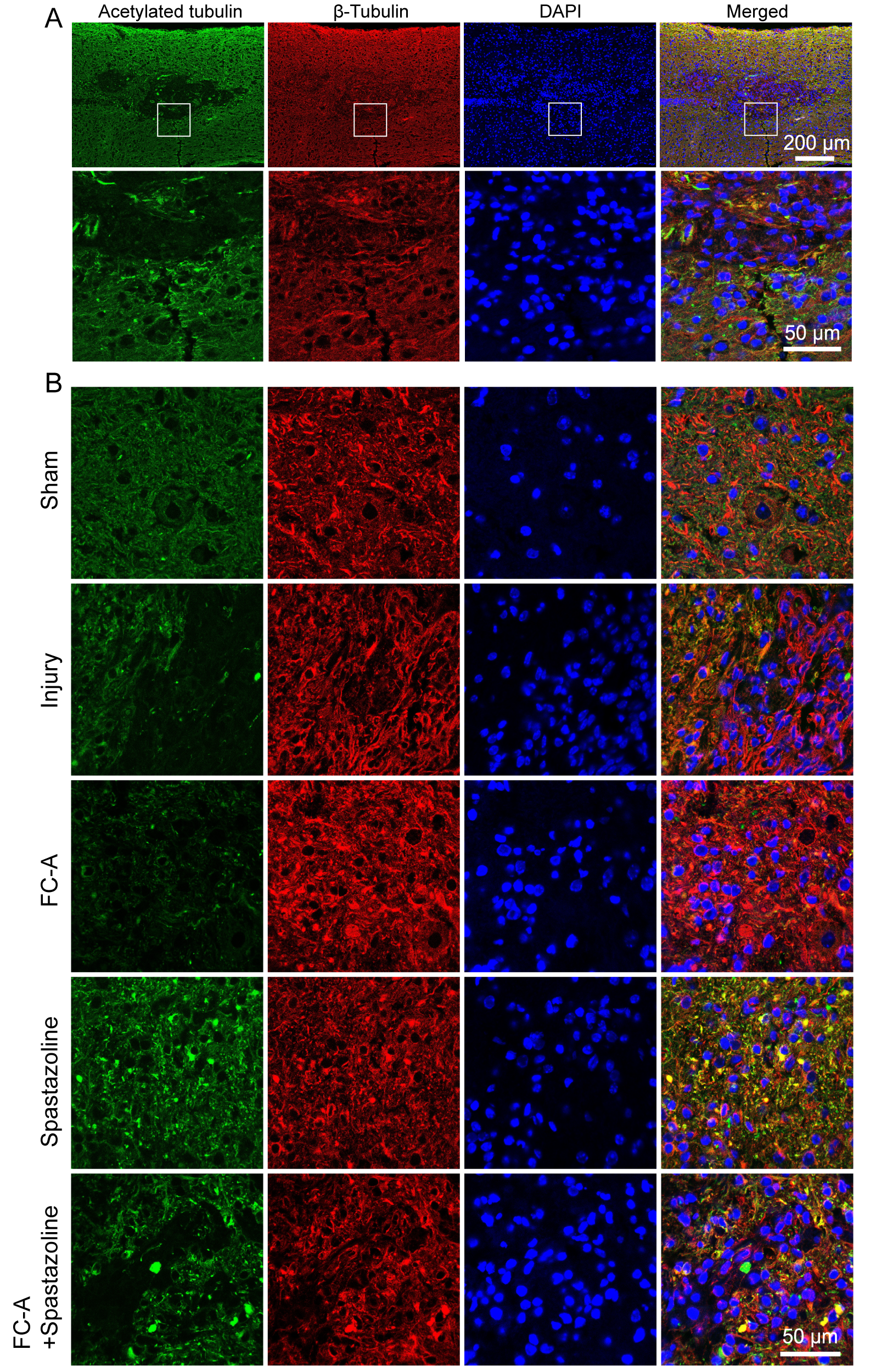

### S9C.jpg

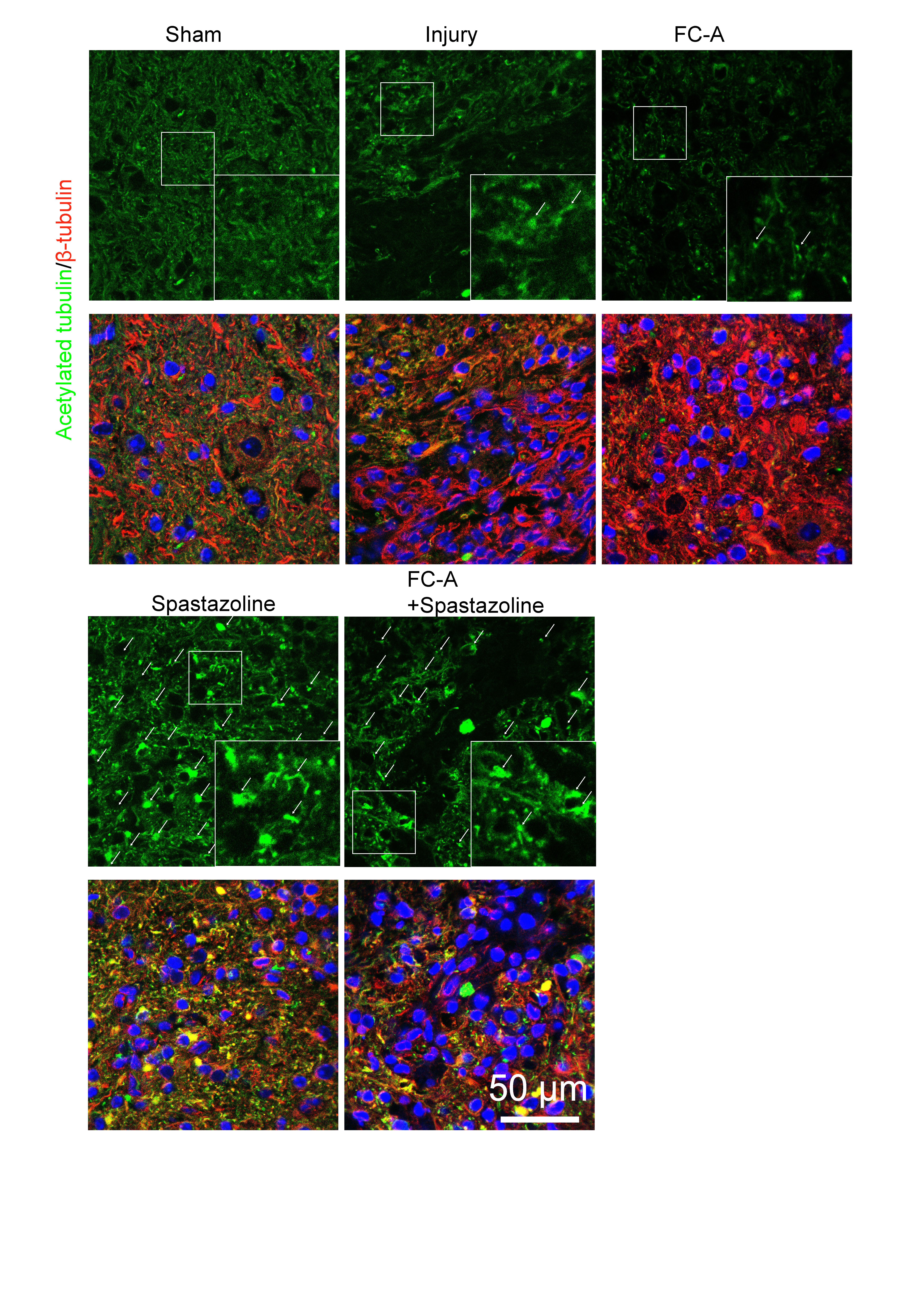

### S10.tif

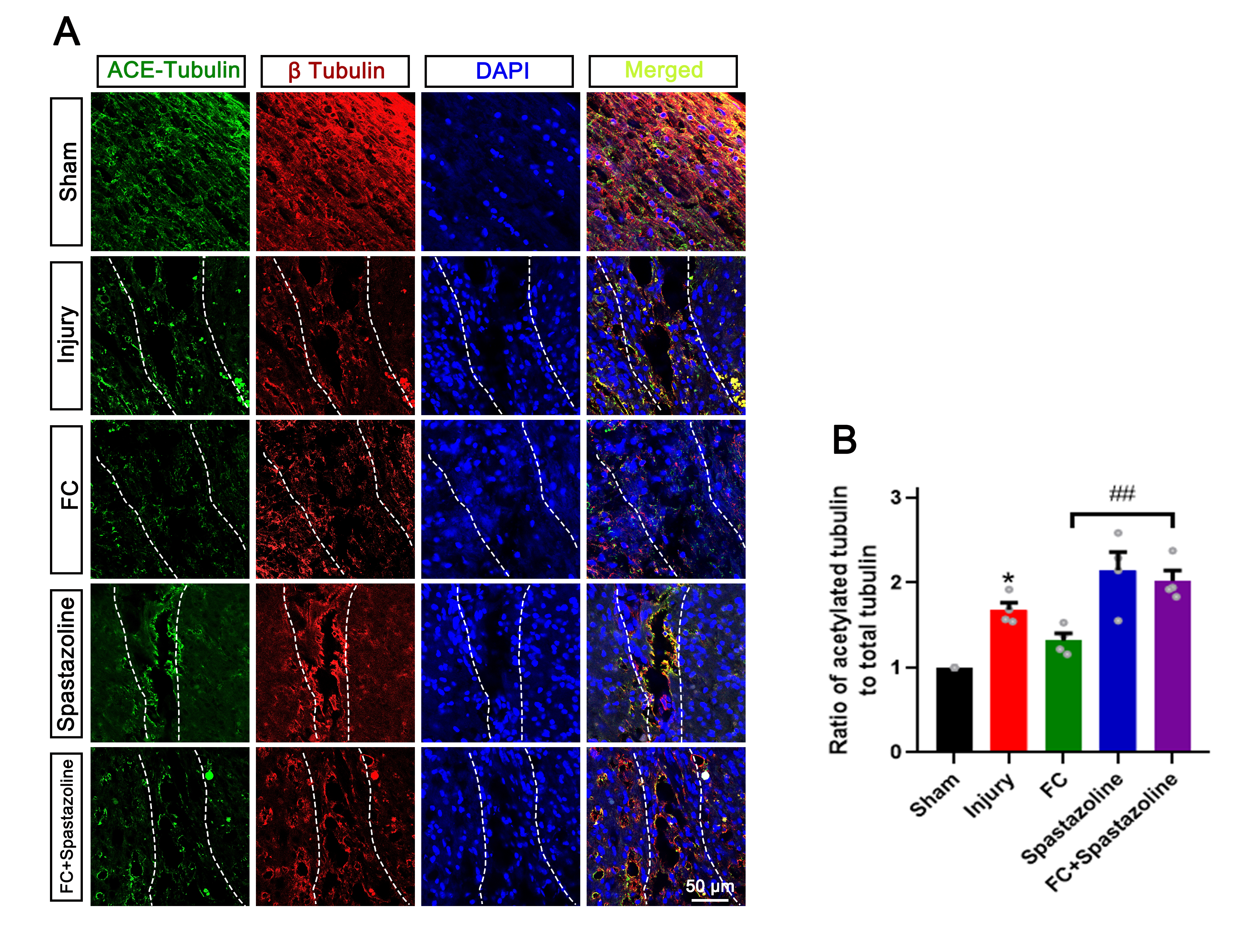

### S11.tif

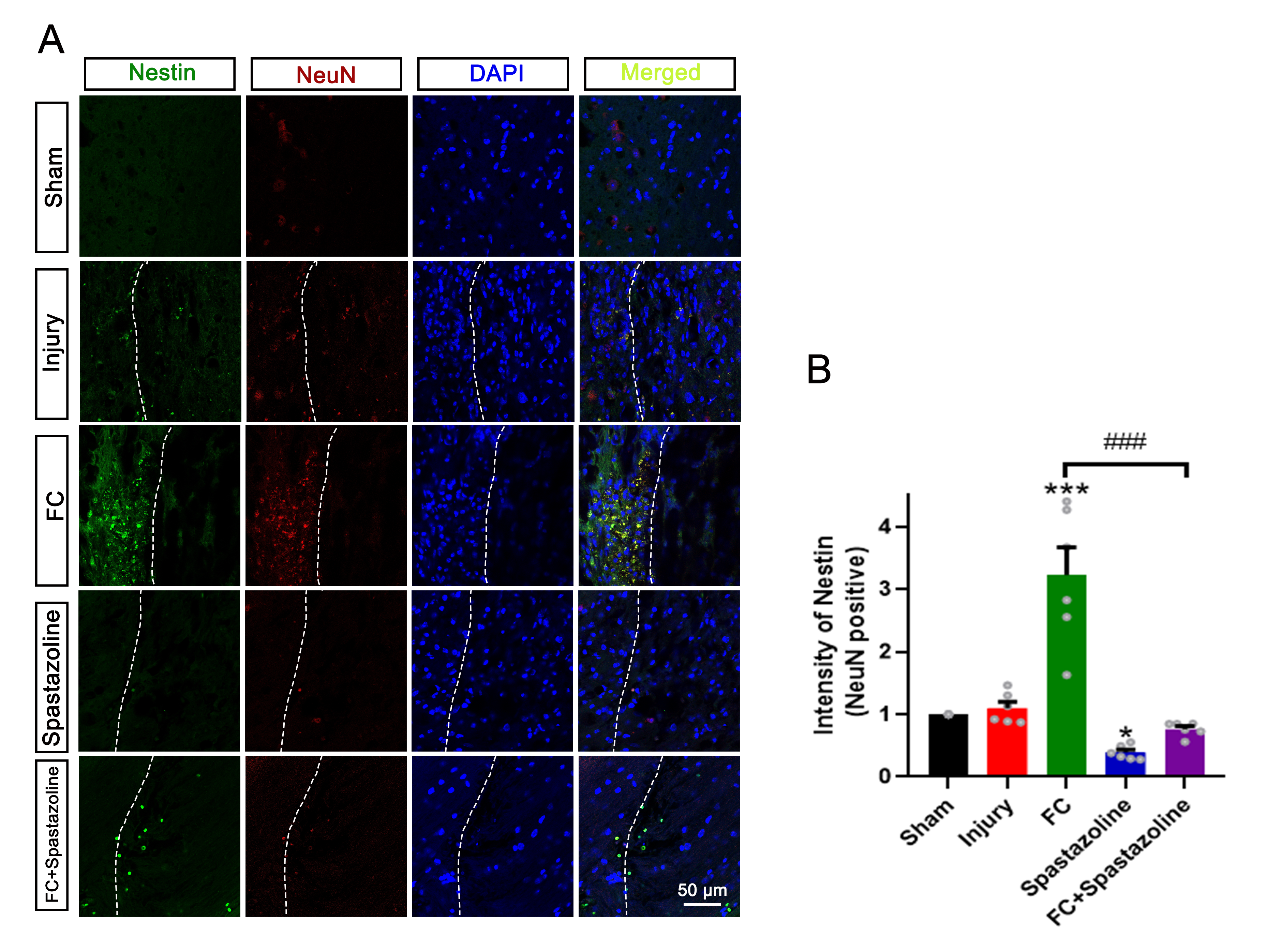

### S13.tif

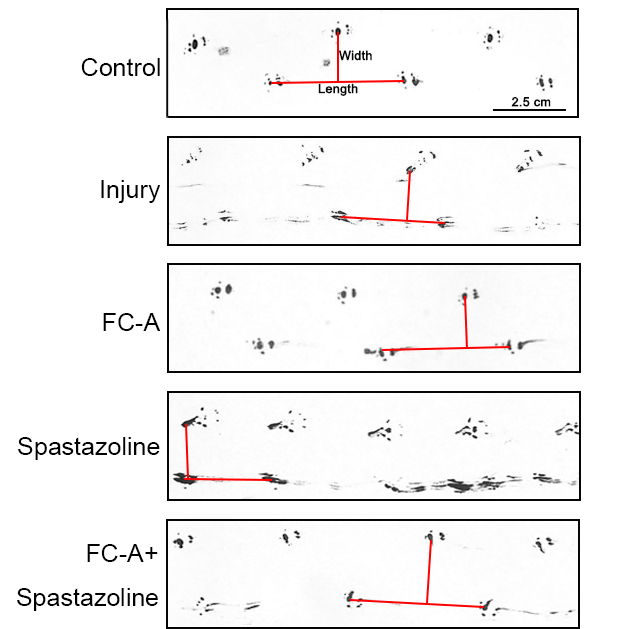
